# Thalamocortical axons control the cytoarchitecture of neocortical layers by area-specific supply of secretory proteins

**DOI:** 10.1101/2021.02.14.431161

**Authors:** Haruka Sato, Jun Hatakeyama, Takuji Iwasato, Kimi Araki, Nobuhiko Yamamoto, Kenji Shimamura

## Abstract

Neuronal abundance and thickness of each cortical layer is specific to each area, but how this fundamental feature arises during development remains poorly understood. While some of area-specific features are controlled by intrinsic cues such as morphogens and transcription factors, the exact influence and mechanisms of action by extrinsic cues, in particular the thalamic axons, have not been fully established. Here we identify a thalamus-derived factor, VGF, which is indispensable for thalamocortical axons to maintain the proper amount of layer 4 neurons in the mouse sensory cortices. This process is prerequisite for further maturation of the primary somatosensory area, such as barrel field formation instructed by a neuronal activity-dependent mechanism. Our results also provide an insight into regionalization of brain in that highly site-specific axon projection anterogradely confers further regional complexity upon the target field through locally secreting signaling molecules from axon terminals.

## Introduction

Making differences within a seemingly uniform entity of cells is a fundamental process observed at various situations in development. The molecular mechanisms that underlie those processes are of great interest for not only developmental biologists but also researchers in stem cell biology and regenerative medicine. While a huge variety of systems or mechanisms have been investigated, providing the basic concepts or principles underlying this issue, there must be yet unidentified mechanisms. The adult mammalian neocortex is entirely composed of six layers of neurons, yet the laminar structure is not uniform throughout the neocortex; the thickness and cellular composition of the layers differ among cortical areas (Brodmann, 1910). For instance, in the sensory cortex, layer 4, which is the main recipient layer of sensory information from the thalamus, is thick and dense, whereas in the motor area, it is thinner. These features are considered to be crucial for proper functions of the cortical areas, as minute differences in these constituents have been shown to be associated with cognitive, mental or psychiatric disorders (Bruining et al., 2015; Reavis et al., 2017; Selten et al., 2018). The developmental mechanisms that regulate the formation of the regionally distinct laminar architecture therefore has long been a major issue in developmental neurobiology.

Previous studies have shown that both mechanisms intrinsic to and extrinsic to the cortex play roles in the formation of cortical areas (reviewed in Cadwell et al., 2019). For instance the secreting factors, emanating from the signaling centers, set up the areal pattern of the neocortex, called cortical area map, through regulating the expression of transcription factors in the cortical primordium (Armentano et al., 2007; Bishop et al., 2000; Fukuchi-Shimogori and Grove, 2001; O’Leary and Sahara, 2008). While the extrinsic mechanisms are less understood, especially at the molecular level, laminar differences among cortical regions correlate well with thalamocortical axon (TCA) projection patterns: abundant TCAs project to sensory areas, which exhibit a thick layer 4, whereas only few axons project to the motor area, which has a thin layer 4. This correlation raises the possibility that TCAs may extrinsically regulate the area differences. In fact, roles of TCAs in cortical development have been investigated particularly with respect to neuronal activity-dependent mechanisms that regulate a histological and functional feature called barrel in the primary somatosensory area (S1) in rodents and ocular dominance column in the primary visual area (V1) of certain species including cat and monkey (Penn and Shatz, 1999; Katz and Crowley, 2002; Gaspar and Renier, 2018). Recent studies demonstrated that TCAs also play instructive roles in the specification of area properties of somatosensory and visual cortices represented by expression of area markers (Chou et al., 2013; Pouchelon et al., 2014; Vue et al., 2013) as well as layer markers, and morphogenesis of cortical neurons (Li et al., 2013; Zhou et al., 2010). Although molecular mechanisms underlying those processes remain uncovered, it was reported that prenatal thalamic neuronal activities and propagation of calcium waves regulate cortical maps prior to sensory processing (Anton-Bolanos et al., 2019; Moreno-Juan et al., 2017). Several studies have also shown that TCA innervation influences neurogenesis in the embryonic cortex. For example, ephrin A5 expressed in TCAs regulates the generation of proper types of cortical progenitor cells and thus neuronal output for cortical layers (Gerstmann et al., 2015). Wnt3 secreted by TCAs controls neuronal differentiation in the cortex at the translational level (Kraushar et al., 2015). However, whether TCAs regulate cytoarchitectural aspects of the cortical layers such as the number of cortical neurons and layer thickness prior to the formation of neuronal connections is not clear. Only one study suggested the involvement of TCAs in cortical laminar formation by showing that thalamic ablation by electrolytic lesion led to alterations in the cortical laminar configuration (Windrem and Finlay, 1991). Yet, it is technically difficult to target thalamic nuclei specifically and reproducibly by surgical manipulation, and the molecular mechanisms by which TCAs control cortical laminar organization have not been uncovered.

To identify TCA-derived extrinsic factors, we previously conducted a screening for thalamus-specific genes by comparing expression profiles of the thalamus and the cortex (Sato et al., 2012). As a result, two genes encoding neuritin 1 (NRN1) and VGF nerve growth factor inducible (Vgf) were found to be expressed specifically in sensory thalamic nuclei including the ventrobasal nucleus (VB). While their mRNAs are not expressed in the cortex, their proteins are detected in cortical layer 4, suggesting that they are transported to the cortex through TCAs. We further found that NRN1 and VGF promoted survival and dendrite growth of cortical neurons *in vitro* (Sato et al., 2012). Although these extrinsic factors are likely to contribute to cortical development *in vivo*, this remains to be validated.

In this study, we investigated the effect of loss of TCAs on neocortical development using transgenic mice, in which thalamic neurons were eliminated postnatally. We employed a diphtheria toxin (DT) receptor (DTR)-mediated cell ablation system combined with a thalamus-specific Cre transgenic mouse line (Arakawa et al., 2014; Buch et al., 2005). As a result, the laminar structure was altered specifically in layer 4, which exhibited marked reduction in the number of neurons in the primary somatosensory cortex. Moreover, TCA-derived factor VGF is necessary for the proper amount of layer 4 neurons in S1 and V1. Interestingly, the barrel organization was impaired in *Vgf*-knockout mice, despite the presence of TCAs and their activities, suggesting that the VGF-dependent quantity control is crucial for the proper barrel field development in cooperation with the activity-dependent process.

## Results

### Distinctions in laminar configuration develop postnatally

We first determined when the laminar differences emerges during mouse cortical development by analyzing the expression of RAR-related orphan receptor beta (RORβ), a layer 4 marker, across the cortical areas from embryonic to postnatal stages using an anti-RORC antibody (see Materials and Methods). At E16 when the production of layer 4 neurons is completed, RORβ expression was high in the anterior and low in the posterior regions (Figure 1A, in 3 mice). RORβ expression then became relatively uniform throughout the cortex, with slight fluctuation in intensity at birth (Figure 1B, in 3 mice). By P7, the staining intensity and thickness of the RORβ-expressing cell layer became greater in the primary somatosensory cortex (S1; Figure 1C, in 5 mice). These data indicated that area-specific laminar characteristics are formed postnatally.

**Figure 1.**
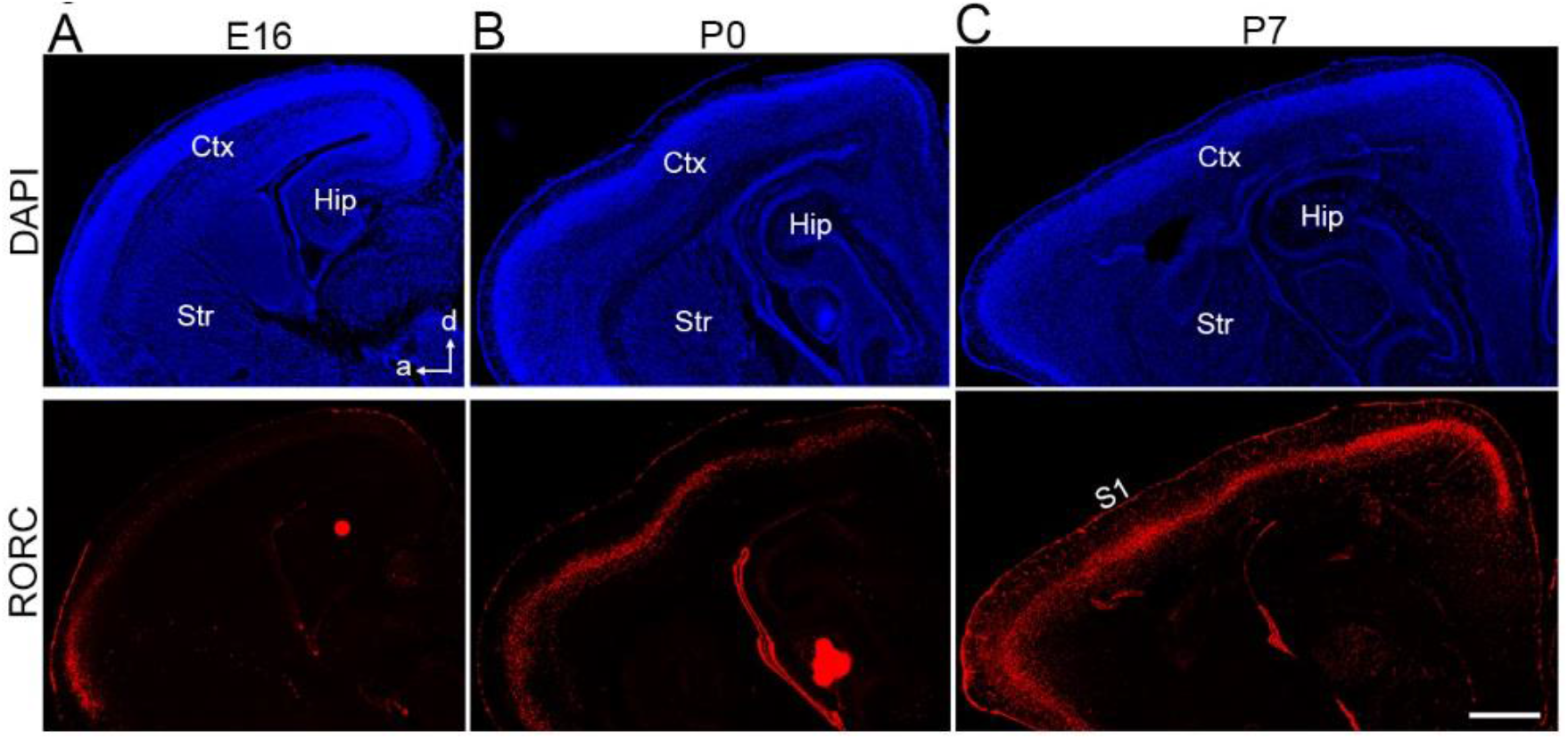
Postnatal emergence of laminar configuration in the cortical area. RORC immunohistochemistry of sagittal sections of E16 (A), P0 (B), and P7 (C) cortices to reveal expression of the layer 4 marker RORβ. Sections are counterstained with DAPI. Ctx, cortex; Hip, hippocampus; Str, striatum; a, anterior; d, dorsal; S1, primary somatosensory area. Scale bars, 500 μm.

Next we asked which mechanisms regulate this transition. As neurogenesis in the cortex is nearly completed at the time of birth (Kwan et al., 2012), it is unlikely that layer 4 neurons are additively generated in S1. Indeed, when EdU was administrated every day from P0 to P4, to label newly generated cells during this period, EdU-labeled cells in layer 4 were all negative for NeuN at P6 (Figure 1-figure supplement 1, in 6 sections from 2 mice), indicating that layer 4 neurons were not newly generated during this period. Thus we concluded that neurogenesis does not take place to form cortical layers in postnatal stages. On the other hand, TCAs innervate the cortex and target layer 4 during the postnatal stages, temporally correlating with the development of the laminar configuration.

### Neonatal ablation of TCAs projecting to S1

To explore the role of TCA projection in layer formation, sensory thalamic neurons were ablated using the Cre-inducible Diphtheria toxin receptor (DTR) mouse system (Buch et al., 2005). A 5HTT-Cre transgenic mouse, in which Cre recombinase activity is detected in sensory thalamic neurons by crossing with an R26-EYFP reporter mouse (Srinivas et al., 2001) (Figure 2A, in 5 mice), was crossed with an R26-DTR mouse (Figure 2B). Diphtheria toxin (DT) was then administrated at P0, and the brains were analyzed at P5-7. At P5, we observed numerous dying cells, detected by ssDNA staining (data not shown) and accumulation of Iba1-immunoreactive microglial cells, which scavenge dead cells, in the thalamus (Figure 2C, in 6 sections from 6 mice), consistent with the previous study (Arakawa et al., 2014). Thalamic cell ablation was further evaluated by immunohistochemistry with an anti-RORC antibody. The VB was dramatically degenerated at P5, and RORα-expressing thalamic neurons were considerably decreased in the VB where Cre expression was the highest among the thalamic nuclei in 5HTT-Cre mice (Figure 2D, in 5 mice). On the other hand, the size of the dorsal lateral geniculate nucleus (dLGN) was not obviously reduced, and RORα-expressing cells were detected. Consequently, the terminals of TCAs were severely diminished in S1 as revealed by 5HTT immunostaining (Figure 2E). Hereafter, we refer to these animals as TCA-ablated mice and animals with DT administration, but without 5HTT-Cre allele, are referred to as control mice.

**Figure 2.**
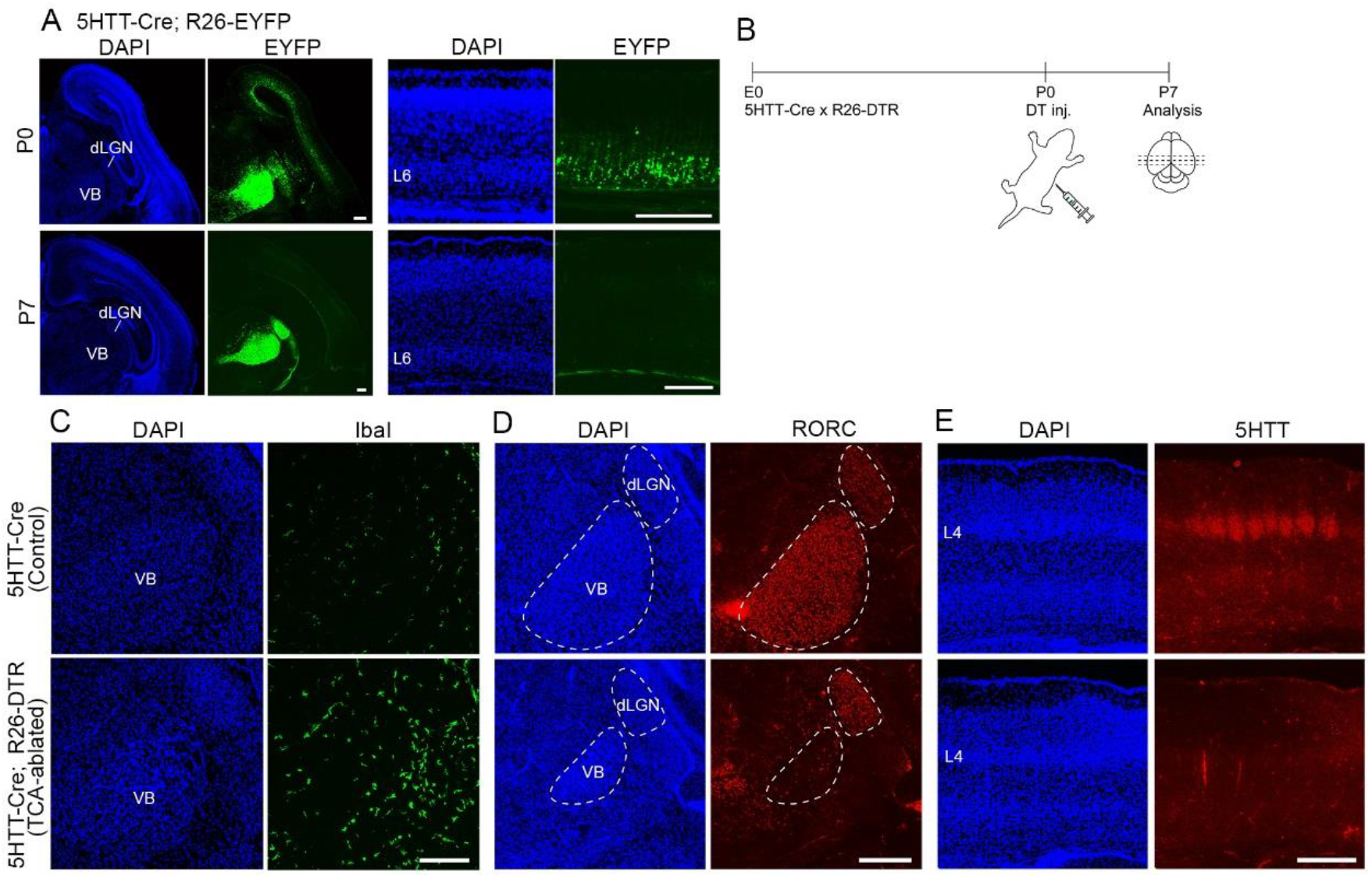
Elimination of TCAs by toxin-mediated thalamic cell ablation *in vivo*. (A) Cre recombinase activities in cross sections of forebrains of P0 (upper panels) and P7 (lower panels) 5HTT-Cre mice. The 5HTT-Cre line was crossed with the R26-EYFP reporter line to allow detection of Cre activity by EYFP fluorescence. EYFP was the most strongly expressed in the VB nucleus and to a lesser extent in the dLGN in the thalamus at P0 (upper left panels), and also weakly in layer 6 (L6) in the cortex (upper right panels), which was lost by P7 (lower right panels). (B) Experimental scheme for TCA ablation. Neonatal pups from a cross of a 5HTT-Cre mouse with a R26-Diphtheria toxin receptor (DTR) mouse were administrated DT intraperitoneally. (C, D) Cross sections of P5 thalamus stained for Iba1 (C) and RORα (D) reveal microglial cells and thalamic nuclei, respectively. Ablation of thalamic neurons was confirmed by loss of RORα expression, especially in the VB. DAPI-stained cell nuclei are packed more densely in the smaller VB than in the control. (E) 5HTT immunohistochemistry of P7 cortices. The amount of TCAs stained for 5HTT was greatly reduced in S1, the target of VB neurons in the TCA-ablated cortex. Note that the 5HTT-positive barrel structure is completely absent in TCA-ablated mice. dLGN, dorsal lateral geniculate nucleus; L4, layer 4; VB, ventrobasal nucleus. Scale bars, 200 μm (A, E), 500 μm (C, D).

To further examine whether TCAs are eliminated or neural connections are altered in TCA-ablated mice, the lipophilic dyes DiA and DiI were injected into two different TCA target areas, S1 and the primary visual cortex (V1), respectively, to label TCAs and thalamic neurons retrogradely (Figure 2-figure supplement 1A, in 4 sections from 2 mice); S1 and V1 are the major target of VB and dLGN, respectively. In control animals, a large number of DiA-labeled thalamic neurons were detected in the VB and the posterior nucleus (PO) (Figure 2-figure supplement 1B, 4 sections from 2 mice). In contrast, in TCA-ablated mice, the retrogradely labeled VB was markedly reduced in size, but the PO was broadly and intensely labeled (Figure 2-figure supplememt 1B, 4 sections from 2 mice). On the other hand, DiI-labeled neurons that project to V1 were found in the dLGN and lateral posterior nucleus (LP) in both control and TCA-ablated mice (Figure 2-figure supplement 1A, 1B, 4 sections from 2 mice for each), and the amount of retrogradely labeled cells were similar in the two groups. Moreover, the fact that DiA-labeled S1-projecting cells were not detected in the dLGN (Figure 2-figure supplement 1B, 4 sections from 2 mice) and medial geniculate nucleus (MG) (Figure 2-figure supplement 1C, 4 sections from 2 mice) suggests that TCA rewiring from the dLGN or MG to S1 did not occur in TCA-ablated mice, unlike previously reported findings (Mezzera and López-Bendito, 2016). Thus, S1 in TCA-ablated mice receives very few axonal inputs from the VB.

### The number of layer 4 cells in S1 is reduced in TCA-ablated mice

To examine impacts of TCA ablation on the laminar structure, the number of cells in each layer was analyzed in control and TCA-ablated cortex using layer makers. RORC immunostaining revealed that layer 4 of S1 was thinner in TCA-ablated than in control mice, leading to poor demarcation of S1 (Figure 3A, 3B, in 5 sections from 3 mice for each). Quantitative analysis revealed that the number of RORβ-expressing cells was decreased to 67% as compared with control cortex (Figure 3C, 15 sections from 7 mice). However, such reduction in layer 4 cells was not observed in non-target areas of VB axons, M1 and V1. V1 still received TCA projection from the remaining dLGN neurons in the experimental situation as described above (Figure 3C, V1, 8 sections from 5 mice; M1, 4 sections from 3 mice; see Figure 2D). Concomitantly, the thickness of the layer abundant in RORβ-expressing cells was significantly reduced in TCA-ablated mice (73.7%, Figure 3D, 16 sections from 7 mice).

**Figure 3.**
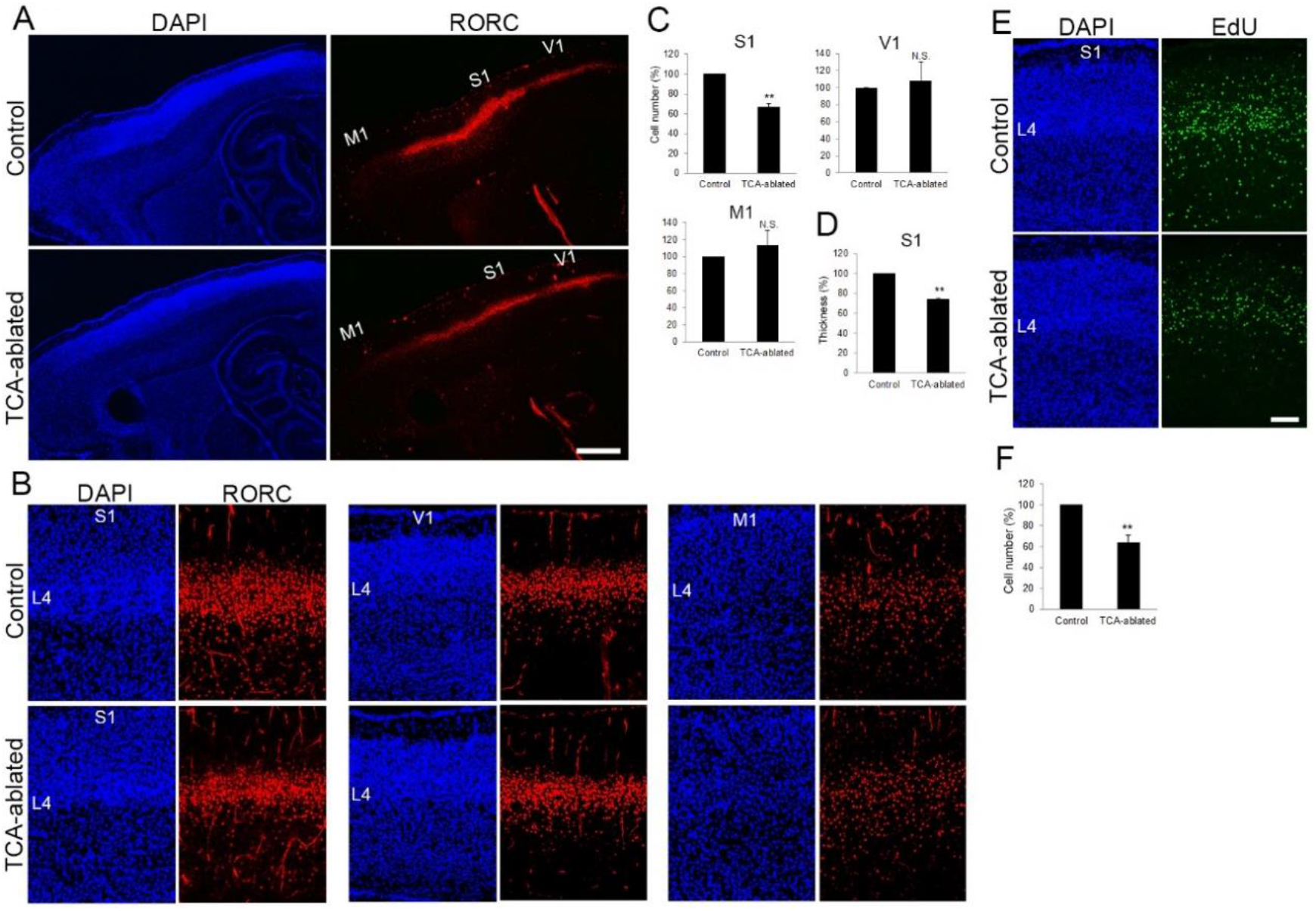
The number of layer 4 neurons is reduced in S1. (A) Sagittal sections of P7 cortices derived from TCA-ablated mice stained with RORC antibody. Expression of the layer 4 marker RORβ was declined, specifically in S1, resulting in poorly demarcated borders between adjacent areas. (B) Coronal section of the cortex of a TCA-ablated mouse at P7, showing RORβ-expressing layer 4 (L4) neurons in S1. (C) Quantification of RORβ expression. RORβ-expressing cells within an 850 μm-wide strip of the cortical wall were counted. Note that the number of RORβ-positive cells was less in S1, but was not changed in V1 and M1, in TCA-ablated mice. Data are presented as a percentage of the expression in control mice (mean ± SEM): S1, 66.75 ± 3.30%, n = 7 mice, *P*=0.0000279; V1, 108.28 ± 22.00%, n = 5 mice, *P*=0.264; M1, 113.24 ± 17.55%, n = 3 mice, *P*=0.362; *t*-test, \*\*\**P* < 0.001. The same numbers of control and experimental animals were used. (D) Thickness of RORβ-expressing layer was also reduced in TCA-ablated mice. Data are presented as a percentage of the expression in control mice (mean ± SEM): 73.69 ± 1.83%, n = 7 mice, *P*=0.00000360; *t*-test, \*\*\**P* < 0.001. (E) Cross sections of S1 cortices of control and TCA-ablated mice at P7, showing the distribution of EdU-positive cells. EdU was injected at E14.7. (F) Quantification of the results. The number of EdU-positive cells within an 850 μm-wide strip of the cortical wall relative to control is shown as the mean ± SEM: 64.37 ± 6.77%, *P*=0.00312; *t*-test, ** *P* < 0.01, n = 5 animals for both control and TCA-ablated mice. L4, layer 4; M1, motor area; S1, primary somatosensory area; V1, primary visual area. Scale bars, 500 μm (A), 100 μm (B, E).

To examine whether the reduction in RORβ-expressing cells in TCA-ablated mice was due to a reduction in cells that had been destined for layer 4, we labeled postmitotic neurons by injecting EdU into the mother at E14.7, when most layer 4 neurons are produced. As expected, EdU-positive cells were preferentially distributed in layer 4 at P7 (Figure 3E, in 12 sections from 5 mice). However, the total number of EdU-positive cells was significantly decreased in TCA-ablated S1 (64% of that in the control; Figure 3E, 3F, 12 sections from 5 mice for each). This reduction was comparable to that in RORβ-expressing cells (Figure 3C). In fact, the proportion of RORβ-expressing cells in EdU-positive cells was not markedly different between TCA-ablated and control mice (Figure 3-figure supplement 1A, 1B, 6 sections from 3 mice for each), indicating that the remaining neurons born at E14.7 maintain RORβ expression in the absence of TCA. Moreover, there was no increase in EdU-positive cells in other layers (Figure 3E, Figure 3-figure supplement 1A, 6 sections from 3 mice). These results suggested that the absolute number of layer 4 neurons decreased in the TCA-ablated S1, and argued against the possibilities of altered RORβ expression or fate change of the layer 4-destined cells to those of other layers.

Next, we examined whether cell death was involved in the reduction in layer 4 cells in S1. Although we used several cell death detection methods (i.e., ssDNA, cleaved caspase 3, Iba1, mRNA of *Bax*, *Bad*, and *Bak*, and DAPI), we could not obtain convincing evidence for significant cell death induction in layer 4 upon TCA ablation (Figure 3-figure supplement 1C, 1D, 8 sections from 7 mice for P2-7; 6 sections from 5 mice for P2-5; 4 sections from 3 mice for P2-4). Given the technical difficulties in detecting dead or dying cells in postnatal compared with adult brain due to the rapid clearance of dead cells (Gohlke et al., 2004; Nagasaka et al., 2010; Wong et al., 2018), it is still possible that cell death is involved in the reduction of layer 4 in TCA-ablated mice.

As Cre recombinase is active not only in the thalamus, but also in cortical layer 6 at P0 (Figure 2A, 10 secions from 5 mice) and the raphe nucleus (data not shown; Arakawa et al., 2014) in 5HTT-Cre mice, we cannot exclude that the layer 4 reduction in 5HTT-Cre; R26-DTR mice is due to ablation of these brain parts rather than the VB in the thalamus. To verify that TCA ablation was indeed responsible for the laminar phenotype of 5HTT-Cre; R26-DTR mice, we ablated VB neurons using a different approach in which a DTR expression plasmid was electroporated into the embryonic dorsal thalamus *in utero* at E11.5, when VB neurons are generated (Figure 3-figure supplement 2A, 2B, 5 sections from 5 mice). Despite variable transfection efficiency in the thalamus, DT administration at P0 induced massive cell death in the VB (Figure 3-figure supplement 2C, 4 sections from 2 mice) and led to a reduction in cell density in the VB (Figure 3-figure supplement 2D, 3 sections from 3 mice). Moreover, 5HTT-positive axon terminals in the cortex were severely reduced (Figure 3-figure supplement 2E, 5 sections from 3 mice), and the number of layer 4 neurons in S1 was decreased (Figure 3-figure supplement 2F, 2G, 4 sections from 3 mice for each), reminiscent of the 5HTT-Cre; R26-DTR phenotype (Figure 3B). Taken together, these results strongly supported the notion that the reduction in layer 4 cells was caused by the loss of TCAs.

### Layers 2/3 and 5 are intact in TCA-ablated mice

To examine whether other layers were also affected by TCA ablation, expression of Brn2 (layers 2/3), Ctip2 (layer 5), and Tbr1 (layer 6) was analyzed. The number of Brn2-positive cells and the thickness of the positive cell layer were not markedly changed in TCA-ablated cortex (Figure 4A, 4D, 4E, 17 sections from 6 mice for each). Similarly, the number and layer thickness of Ctip2-expressing cells were not greatly affected (Figure 4B, 4D, 4E, 13 sections from 3 mice for each). In contrast, Tbr1-expressing cells were partially reduced in TCA-ablated mice (Figure 4C, 4D, 6 sections from 3 mice for each), likely due to DT-induced cell death of the layer 6 neurons that express Cre recombinase at P0 as mentioned above (Figure 2A). Indeed, we observed signs of cell death in layer 6 (i.e., ssDNA-positive, accumulation of microglia; data not shown). However, the thickness of layer 6 was not markedly changed (Figure 4C, 4E, 6 sections from 3 mice for each). Collectively, these results supported the specificity of the effects of TCA ablation on cortical layer formation in that the effect was restricted to the target layer of TCA projection.

**Figure 4.**
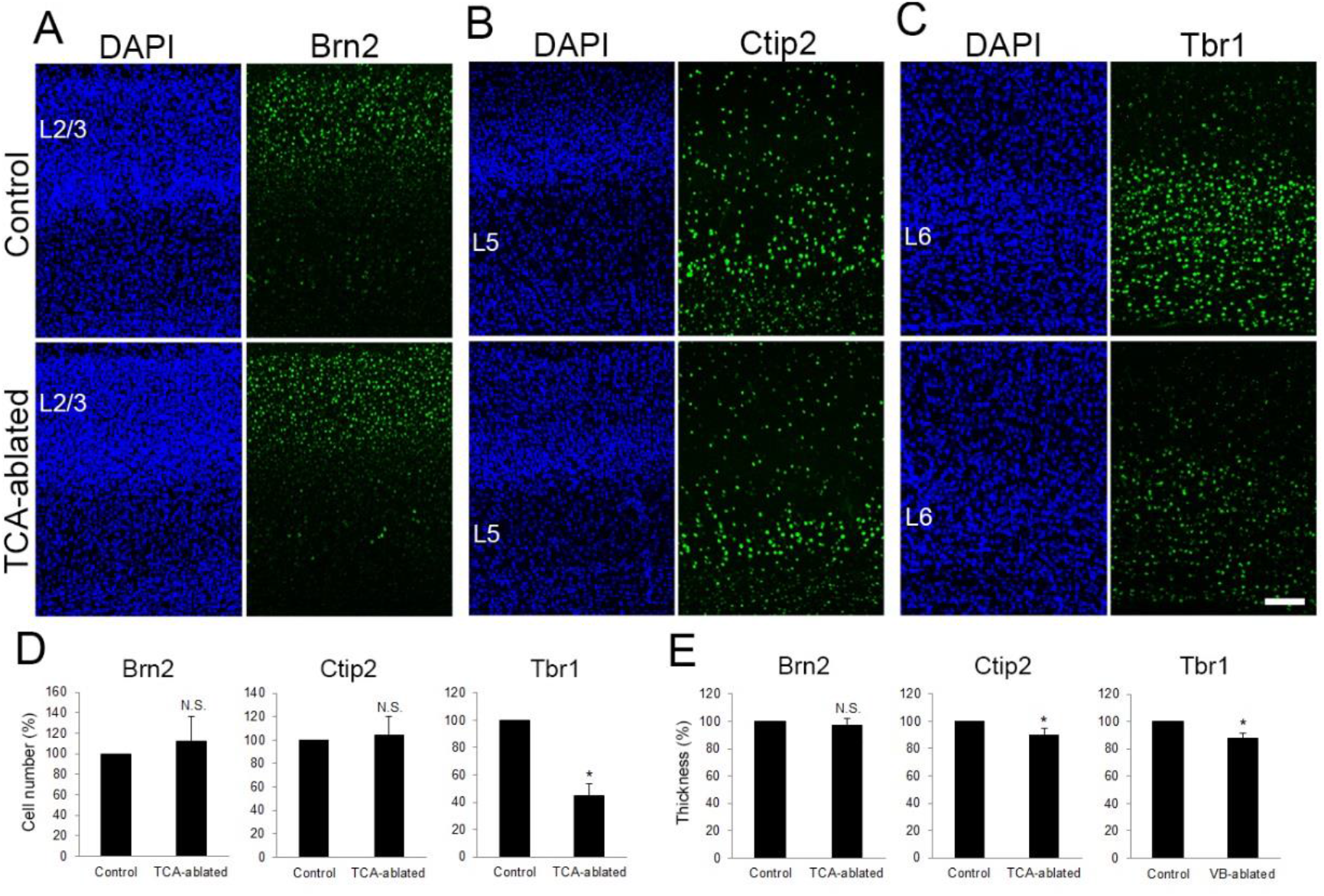
Layers 2/3 and 5 appear intact upon TCA elimination. Cross sections of S1 cortices of TCA-ablated mice at P7 stained for Brn2 (layer 2/3)(A), Ctip2 (layer 5)(B), and Tbr1 (layer 6)(C). Expression of Brn2 and Ctip2 were not markedly changed, whereas Tbr1-positive cells were decreased in TCA-ablated mice. (D, E) Quantitative analyses of the number of cells (D) and layer thickness (E) of TCA-ablated S1 relative to the control. Cells positive for each marker within an 850 μm-wide strip of the cortical wall were counted. Data are presented as a percentage of the control (mean ± SEM): Brn2, 112.70 ± 23.79%, n = 6 animals, *P*=0.308; Ctip2, 104.56 ± 15.86%, n = 3 animals, *P*=0.400; Tbr1, 44.50 ± 23.79%, n = 3 animals, *P*=0.0118; *t*-test, \**P*<0.05. The same numbers of control and experimental animals were used. (E) Thickness of the layers where Brn2-, Ctip2-, and Tbr1-positive cells were densely distributed. Cell numbers are presented relative to the control (mean ± SEM): Brn2, 97.40 ± 5.12%, n = 6 mice, *P*=0.316; Ctip2, 90.23 ± 4.38%, n = 5 mice, *P*=0.0447; Tbr1, 87.80 ± 3.43%, n = 3 mice, *P*=0.0354; *t*-test, \**P* < 0.05. Scale bar, 100 μm.

### The number of layer 4 neurons is restored by forced expression of NRN1 and VGF in the cortex of TCA-ablated mice

Regarding the molecular basis of TCA-dependent layer 4 formation, we hypothesized that extracellular molecules emanating from TCA terminals are involved. As our previous study showed that NRN1 and VGF, which promote cortical cell survival and dendritic growth, are localized in TCA terminals (Sato et al., 2012), the expression and action of these molecules were investigated. As expected, the expression of both NRN1 and VGF was lost in the thalamic nuclei and their axon terminals in layer 4 of S1 in TCA-ablated mice (Figure 5-figure supplement 1A, 1B, 2 sections from 2 mice for each). To examine whether these proteins play any role in the reduction in layer 4 neurons in the TCA-ablated S1, we overexpressed these factors in layer 4 cells by *in utero* electroporation prior to TCA ablation (Figure 5A, 5B, 6 mice). As a result, the number of layer 4 cells, which were defined by RORC immunoreactivity, was completely restored to the control level (Figure 5C, 5D, 11 sections from 6 animals for each). Curiously however, we did not observe an additive increase in layer 4 neurons by overexpression of these factors in the presence of TCA (Figure 5-figure supplement 2B, 10 sections from 5 mice). Taken together, this result suggested that NRN1 and VGF as TCA-derived factors function to maintain the layer 4 neuronal number during postnatal neocortical development.

**Figure 5.**
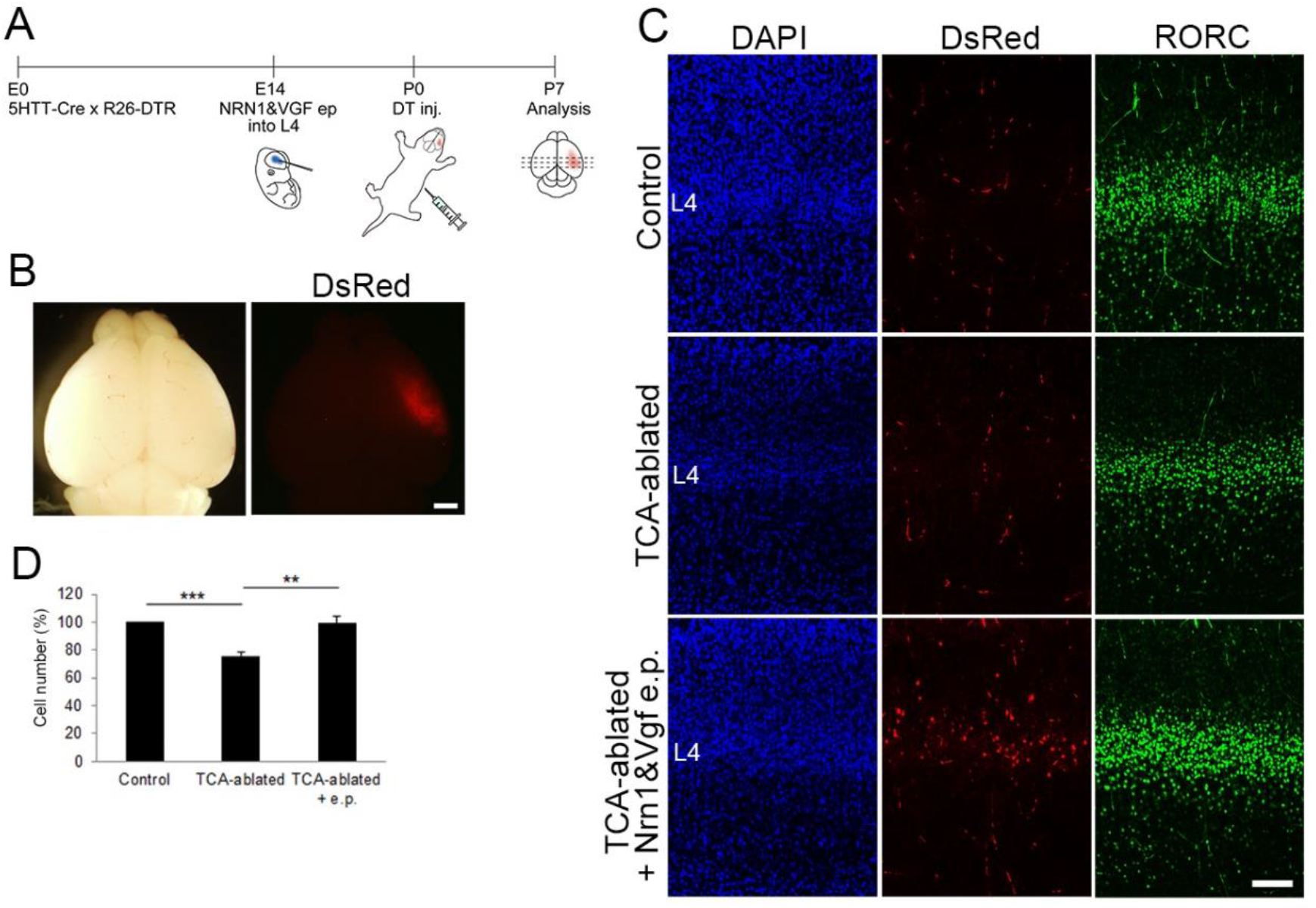
Overexpression of NRN1 and VGF rescues layer 4 formation in TCA-ablated mice. (A) Schematic representation of the experimental procedure. (B) P7 electroporated brain showing DsRed signal in the right hemisphere. (C) Cross sections of S1 cortices of control, TCA-ablated, and TCA-ablated + electroporated (e.p.) mice stained for DsRed and RORβ. (D) Quantitative analysis of RORβ-expressing cells. Data are presented as a percentage of the control (mean ± SEM): TCA-ablated, 75.61 ± 2.87%, n = 6 animals, *P*=0.000186; TCA-ablated+electroporation, 99.52 ± 4.54%, *P*=0.00106; *t*-test, \*\**P* < 0.01, \*\*\**P* < 0.001, n = 6 animals. Scale bars, 1 mm (B), 100 μm (C).

### Genetic inactivation of *Vgf* results in a reduction in layer 4 neurons in S1

We further addressed whether NRN1 and VGF are in fact necessary for the regulation of layer 4 development in S1 by using CRISPR/Cas9-mediated gene editing (Harms et al., 2014). We designed three single-guide RNAs (sgRNAs) cutting exons of *Nrn1* and *Vgf* to induce frame shifts resulting in failure of protein translation of both NRN1 and VGF. By electroporating the sgRNAs and Cas9 protein into fertilized eggs, mutations were induced in the genomic sequences of *Nrn1* and *Vgf* allele near designed sgRNAs. While we could not obtain double-null mice, probably due to lethality during embryonic or early postnatal stages, single-mutant mice for *Nrn1* or *Vgf*, both of which harbored null mutations, were collected at P8. We confirmed the loss of NRN1 and VGF protein by western blotting (data not shown) and immunohistochemistry (Figure 6A, 5 sections from 5 mice; data not shown for NRN1). Immunoreactivity of VGF was completely abolished in the VB in the thalamus, where it is endogenously expressed as described previously (Sato et al., 2012), as well as in layer 4 of S1, where TCAs from the VB terminate. As reported previously (Hahm et al., 1999), the body weight of *Vgf*^−/−^ mice was significantly lower than that of *Vgf*^+/+^ control mice. Although the size of the cortex was slightly larger, gross brain anatomy was not markedly different (Figure 6B, 6C, wild-type, 4 mice; *Vgf*^*−/−*^, 5 mice). Importantly, the thalamic structure was not affected: the VB appeared intact in terms of shape, size, and cell density (Figure 6D, 5 sections from 5 mice). In addition, TCAs revealed by 5HTT immunohistochemistry were present in layer 4 of S1, similar to the wild-type (Figure 6D, 6 sections from 3 mice), indicating that thalamocortical projection was formed properly. The effect of loss of TCA-derived VGF from the cortex was evaluated by RORβ immunohistochemistry. The number of positive cells in layer 4 was markedly reduced in S1 and in V1 (Figure 7A, 7B, 7E, 8 sections from 5 mice for each). Other layers, including layers 2/3 and 5, were not affected except a slight increase in Ctip2-positive layer 5 (Figure 7C, 7D, 7F, Brn1, 20 sections from 5 mice; Ctip2, 18 sections from 5 mice). In spite of the significant decrease in layer 4 cells, layer thickness was more mildly decreased than the TCA ablation (91.3%, Figure 7G, 8 sections from 5 mice; see Figure 3D). These findings suggested that TCAs regulate two aspects of area-specific layer 4 formation separately, in that layer thickness is controlled by the presence of TCAs, and the number of layer 4 neurons is regulated by TCA-derived VGF. To validate the significance of this regulation in the further development of S1, the barrel formation, an unique feature observed in S1 in rodent (Kawasaki, 2015), was examined. The characteristic columnar repetitive arrangement of cells in layer 4 was impaired in *Vgf* KO, such that cells were more evenly distributed along the tangential plane of layers in the mutant S1 revealed by a quantitative image analysis (Figure 7H, 7I, wild-type, 3 sections from 3 mice; *Vgf*^*−/−*^, 4 sections from 4 mice). Consistent with the presence of TCA terminals in *Vgf*-KO layer 4 (see Figure 6D), expression of *Btbd3* in layer 4 neurons, which is known to be dependent on neuronal activities from TCAs (Matsui et al., 2013), was substantially recognized unlike the TCA-ablated cases (Figure 7-figure supplement 1, 4 sections from 2 mice). Therefore, the abnormal barrel organization in *Vgf*-KO is unlikely due to failure of activity-dependent processes, suggesting that the proper neuronal number in layer 4 is prerequisite for this process to operate. In contrast to *Vgf*^−/−^ mice, *Nrn1*^−/−^ mice exhibited a normal layer 4, such that the neuronal number was comparable to that in control *Nrn1*^+/+^ mice (Figure 7-figure supplement 2, 2 sections from 1 mouse). Taken together, these results indicated that VGF regulates of the neuronal number in layer 4 of S1, and that NRN1 is dispensable in this process.

**Figure 6.**
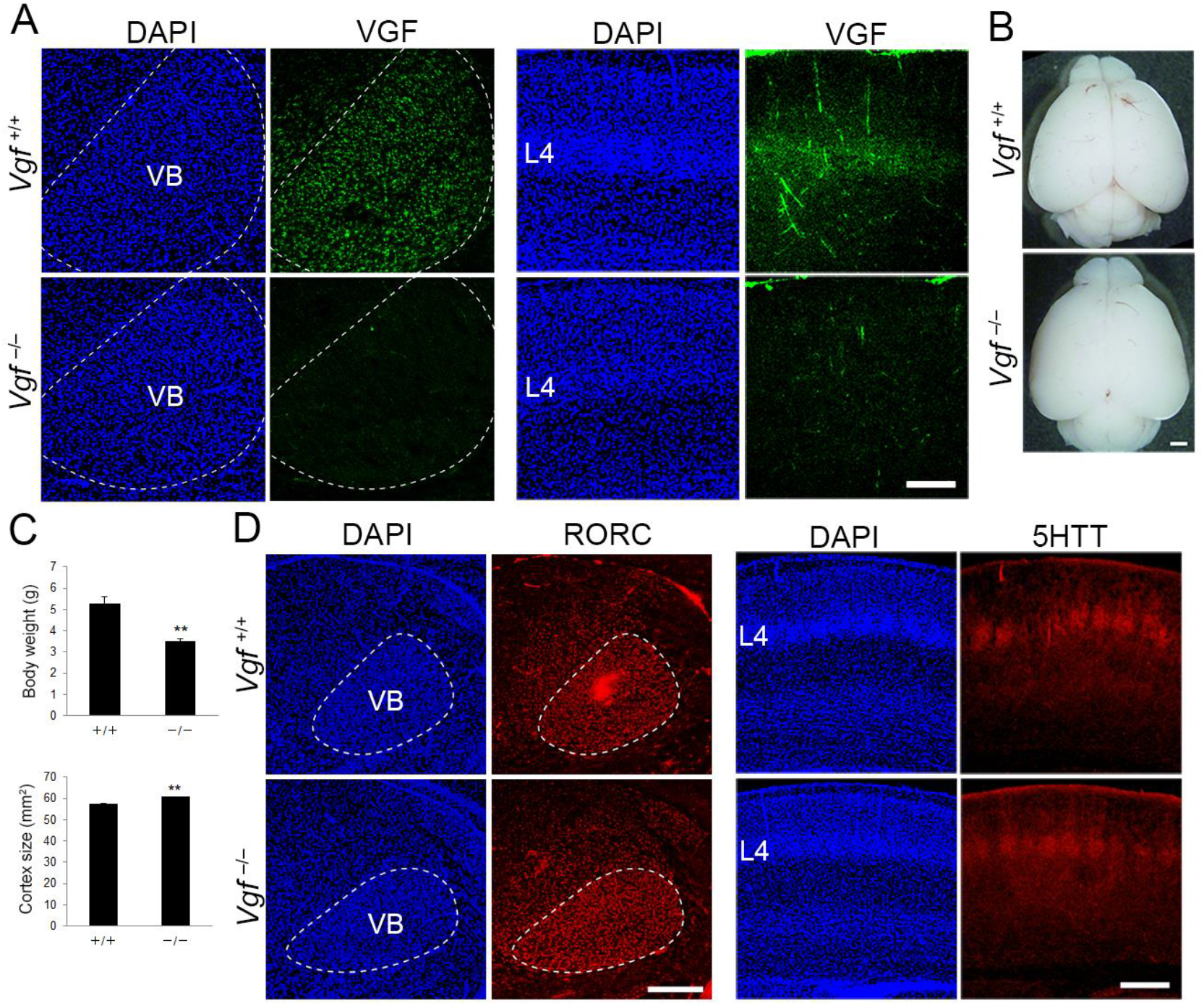
Generation of *Vgf*-KO mice using the CRISPR/Cas9 system. (A, D) Coronal sections of thalamus (left groups) and S1 cortex (right groups) of wild-type and *Vgf*-deficient mice at P8 stained with anti-VGF (A), -RORC (D), and −5HTT (D) antibodies. Note that VGF protein is lost in the VB and cortical layer 4 of *Vgf*^*−/−*^ mice, whereas expression of RORβ in the VB and the presence of 5HTT-positive TCA terminals in cortical layer 4 were not affected in *Vgf*^−/−^ mice. (B) Dissected brains of wild-type and *Vgf*^*−/−*^ mouse at P8. (C) Quantitative data of body weight and cortical surface area presented as the mean ± SEM: body weight: wild-type, 5.29 ± 0.42 g, n = 4 mice; *Vgf*^*−/−*^, 3.51 ± 0.05 g, n = 5 mice; *t*-test, *P*= 0.00317; cortical surface area: wild-type, 57.43 ± 0.62 mm^2^, n = 4 mice; *Vgf*^−/−^, 60.76 ± 0.37 mm^2^, n = 5 mice, *P*= 0.00285; *t*-test, \*\**P* < 0.01,. Scale bars, 100 μm (A), 1 mm (B), 500 μm (C).

**Figure 7.**
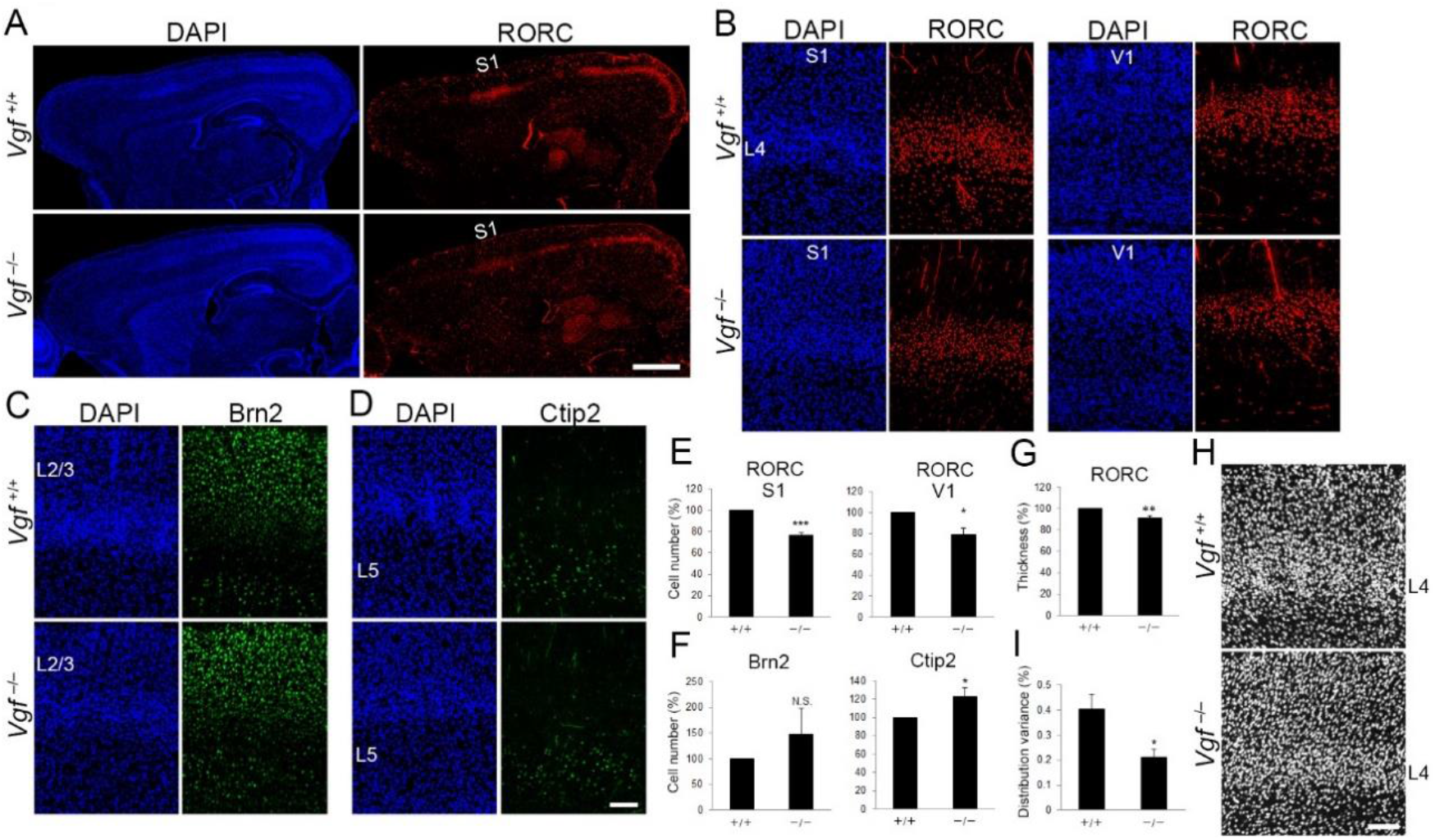
*Vgf*-deficiency causes a reduction in layer 4 neurons in S1. (A) RORC immunostaining of sagittal sections of P7 cortices of wild-type and *Vgf*^−/−^ mice revealed the reduction in layer 4 in the S1 area in the mutant. (B-D) Coronal sections of P8 cortices of wild-type and *Vgf*^−/−^ mice stained with anti-RORC (B), anti-Brn2 (C), and anti-Ctip2 (D) antibodies. (E, F) Quantification of the results. Data are presented as a percentage of the wild-type control (mean ± SEM): RORC S1, 76.72 ± 1.79%, n= 5 *Vgf*^−/−^ mice, *P*=0.000102; RORC V1, 79.25 ± 6.14%, n= 5 *Vgf*^−/−^ mice, *P*= 0.0139; Brn2, 147.57 ± 50.63%, n = 5 *Vgf*^−/−^ mice, *P*=0.200; Ctip2, 122.67 ± 9.93%, , n = 5 *Vgf*^−/−^ mice, *P*=0.0422; *t*-test, \**P* < 0.05, \*\*\**P* < 0.001. Three wild-type mice were used as the reference. (G) The thickness of RORβ-expressing layer was slightly reduced in *Vgf*^−/−^ mice, but not as much as in the case of TCA ablation. Relative thickness to the wild-type was 91.34 ± 2.24%, n = 5 *Vgf*^−/−^ mice, *P*=0.00904; *t*-test, \*\**P* < 0.01. Three wild-type mice were used as the reference. (H) DAPI staining of coronal sections showed barrel structure as uneven and repetitive distribution of cellular nucleus in layer 4 of wild-type, but not of *Vgf*^−/−^ mice. (I) Distribution variance of layer 4 cells as the mean ± SEM: wild-type, 0.403 ± 0.060, n = 3 mice; *Vgf*^*−/−*^, 0.213 ± 0.031, n = 4 mice, *P*=0.0340; *t*-test, \**P* < 0.05. Scale bars, 1 mm (A), 100 μm (B-D, H).

## Discussion

How the laminar configuration is formed in the developing cortex is one of the fundamental questions in developmental neurobiology, and the underlying molecular mechanisms have not yet been fully elucidated. Here, we investigated the influence of TCAs on cortical laminar formation by ablating TCAs from the thalamic VB *in vivo*. We found that the number of layer 4 neurons in S1 was decreased in the absence of TCAs during postnatal stages. This was rescued by overexpression of axon-derived secreted proteins, NRN1 and VGF, in cortical layer 4. Furthermore, genetic disruption of *Vgf* resulted in a reduction in the layer 4 neuronal number. Collectively, these results indicated that TCAs are required for the formation of the area-specific laminar structure by regulating the number of layer 4 neurons through VGF. Our finding that the number of layer 4 neurons was reduced by 34%, which resulted in poorly distinguishable S1 from neighboring areas in TCA-ablated mice, indicates that TCAs play a substantial and major role in specialization of S1, which is characterized by a thick layer 4. Thus, highly site-specific axon projection, which depends on initial regional patterning of the target fields, in turn contributes to generation of the further regional diversity and complexity within the target tissues in a remote fashion.

While the present findings explain why layer 4 is thick in the somatosensory cortex, it is currently unclear whether the same mechanism accounts for the very thin layer 4 in the motor cortex. It would be interesting to test experimentally whether abnormal projection of TCAs to the motor cortex or artificial supply of TCA-derived factors in the motor cortex would thicken layer 4 and increase the neuronal number. While TCA-ablated mice showed almost no defect on V1, likely due to a small number of cell death in dLGN that innervates V1, *Vgf*-KO mice exhibited significant reduction in layer 4 neurons in V1. These observations suggest that the regulation of the neuronal number of layer 4 by TCAs via VGF is a common mechanism operating widely in sensory areas.

The exact cellular events that led to the reduction in layer 4 neurons in the TCA-ablated cortex remain to be determined. Although most likely, specific cell death underlies this phenomenon, we did not detect obvious signs of cell death in this process. Previous studies have analyzed cell death in the postnatal cortex in wild-type mice (Ferrer, 1992; Gohlke et al., 2004; Spreafico et al., 1995; Thomaidou et al., 1997; Verney et al., 2000), however, cell death in specific layers or areas in relation to the area-specific laminar structure has not been reported. Instead, a relatively uniform distribution of dying cells across layers and areas was reported in these studies. We also examined *Bax* and *Casp3* mutant mice; the laminar configuration across the areas appeared to be normal (HS, KS, unpublished observation). These observations suggest that relatively thinner layer 4 in other areas compared to S1 may not depend on specific cell death, yet it is still possible that it plays a role in the absence of TCAs in S1.

Another possibility is neuronal migration. It has been reported that mismigration of layer 4-destined neurons to layers 2/3 upon knockdown of *Protocadherin20* resulted in respecification of these neurons to acquire layer 2/3 characteristics, suggesting a positional influence on cortical neuron fate (Oishi et al., 2016). However, we found no evidence that neurons destined to layer 4 migrated to other layers (Figure 3, 4) and changed their fate in the TCA-ablated S1 (Figure 3-figure supplement 1). Beside radial migration, neurons in layer 4 might migrate tangentially across areas, resulting in the area-specific laminar features in normal development. Such a process might have been impaired by the loss pf TCAs, leading to a failure of accumulation of neurons into layer 4 of S1. To test this possibility, we tracked layer 4 neurons in the frontal cortex using the photo-switchable fluorescent protein kikGR (Nishiyama et al., 2012) in living pups from P0 to P7 by a laser-scanning confocal macroscopy. We did not detect any obvious signs of tangentially migrating layer 4 neurons; the labelled cells were retained within the original area of photoconversion in the motor area (HS and KS, unpublished results). Nevertheless, we might have overlooked cell migration because of limited detection sensitivity. In future, it may be worth pursuing this possibility using more sophisticated and super-sensitive methods recently published, through which the authors successfully observed activity-dependent dynamic dendrite rearrangement during the course of barrel formation in S1 (Mizuno et al., 2014). This method may allow us to observe if layer 4 neurons in S1 disperse into or fail to accumulate from surrounding areas in TCA-ablated mice. Similar technology might also enable us to determine whether cells in layer 4 undergo cell death in the absence of TCAs or their actions.

There are two classical models for the development of cortical areas. One is the protomap model, in which the areal properties are predisposed in the regional identity of the cortical progenitors (Rakic et al., 2009). The other is the protocortex model, in which the cortical primordium is generated essentially homogeneous and is patterned into areas later by cues from the thalamic axons (O’Leary, 1989). In theory, these models are not mutually exclusive (Sur and Rubenstein, 2005), but can be reconciled as serial homology and refinement model recently proposed (Cadwell et al., 2019), in which cortical area development can be divided into serial three steps: 1) protomap, 2) area-specific maturation, and 3) activity-dependent refinement; processes postulated in the protocortex model are involved in 2) and 3). Previous studies on the roles of TCAs in cortical area development have led to the notion that TCAs indeed regulate areal size and characteristic gene expression in the cortex (Chou et al., 2013; Pouchelon et al., 2014; Vue et al., 2013); Moleno-Juan et al., 2017). Moreover, it has been reported that neural activity from TCAs controls neuronal morphology and barrel formation in a later stage (Li et al., 2013; Narboux-Neme et al., 2012). In this context, our present notion can be regarded as a process in 2) area-specific maturation: VGF released from TCA terminals control the number of cortical neurons which is indispensable for the subsequent activity-dependent interaction between thalamic and layer 4 neurons as the barrel organization in S1 was significantly impaired in *Vgf*-KO mice despite the presence of TCAs.

Afferent-derived proteins play important roles in neural development. For example, ephrin A5 and Wnt3 from TCAs regulate differentiation of cortical neurons as mentioned above (Gerstmann et al., 2015; Kraushar et al., 2015). In other nervous systems, afferent-derived proteins regulate neurogenesis and synapse formation in their target region (Huang and Kunes, 1998; Sanes and Lichtman, 2001). In this study, we demonstrated that TCA-derived secretory protein VGF is necessary to maintain a sufficiently high neuronal number in cortical layer 4. VGF is widely expressed in the sensory thalamic nuclei, including the VB, dLGN, and medial geniculate nucleus (Sato et al., 2012), suggesting that this cue is commonly used for higher-order differentiation of sensory cortices. In fact, the number of layer 4 neurons in V1, which is a target of dLGN, was reduced in *Vgf*-KO mice (Figure 7B, ‘E). Although disruption of NRN1, another TCA-derived secretory protein, did not affect the number of layer 4 neurons, it may play roles in other aspects of layer formation, e.g., dendritic growth of spiny stellate neurons, as previously shown *in vitro* (Sato et al., 2012).

Our finding that a TCA-derived factor contributes to areal differentiation of the cerebral cortex may provide an insight into the generation of diversity in the areal pattern across mammalian species. In general, there is a strong correlation between dependence on particular sensory modality adopted for living environments and the proportional development of the corresponding areas (Krubitzer, 2007). In fact, experimental manipulation of the size of thalamic nuclei resulted in appropriate alteration of the areal pattern (Chou et al., 2013; Vue et al., 2013). Moreover thalamic calcium wave plays critical roles in coordinating areal size prior to sensory processing (Moreno-Juan et al., 2017). Thus, mammals seem to have acquired coordinated evolution of thalamic sensory modality components and cortical areal characteristics to support their ecological diversity, resulting in the present prosperity worldwide. In this regard, it would be intriguing to explore the roles of the thalamus-derived factors, such as VGF or NRN1, along with the calcium wave mentioned above, in cortical area diversity among mammalian species. It is worth noting that both NRN1 and VGF are induced by neuronal activities in various brain regions including the cortex, thalamus, and hippocampus in later developmental stages and in adults (Corriveau et al., 1999; Harwell et al., 2005; Snyder et al., 1997). Although it is not known how these factors are induced in the thalamus during area formation, it could be a potential link between evolution of a peripheral sensory system and its corresponding cortical area.

## Acknowledgements

The authors thank J. Kusuura for excellent technical support. We are also grateful to Drs T. Shimogori and Y. Yamaguchi for discussion, reagents, and mutant mice. This work was supported by KAKENHI grants from Japanese government (KM101-2587054400, KM100-2633200, KM101-18K1483900 to HS; 06J08049, 21870030, 24790288, 15K19011, 16H01449, 17H05771 to JH; 18GS0329-01, 16K07375 to KS) and Grant-in-Aid for Scientific Research on Innovative Areas “Platform of Advanced Animal Model Support” from the Ministry of Education, Science, Sports and Culture of Japan. We thank the Liaison Laboratory Research Promotion Center at IMEG, Kumamoto University for technical support.

## Author contributions

HS designed and performed all the experiments and analyses, and JH and KS assisted them. TI generated and provided the 5HTT-Cre line. KA generated *Vgf*- and *Nrn1*-KO mice. HS and NY conducted the initial stage of the research. HS and KS wrote the paper.

## STAR Methods

### Animals

Time pregnant ICR mice (SLC Japan) were used for immunohistochemistry, *in utero* electroporation and EdU administration. The day of vaginal plug detection was designated as embryonic day 0 (E0), and the day of birth as postnatal day 0 (P0). 5HTT-Cre mice were previously generated (Arakawa et al., 2014). R26-EYFP mice were a generous gift from Dr. Constantini (Columbia University, USA) (Srinivas et al., 2001). R26-DTR mice were purchased from The Jackson Laboratory (C57BL/6-*Gt(ROSA)26Sor*^*tm1(HBEGF)Awai*^/J). For cell ablation, 40 ng of DT was administrated once into pups of 5HTT-Cre; R26-DTR mice by intraperitoneal injection through the back skin at P0 (Calbiochem, #322326). All experiments were carried out following the Guidelines for Laboratory Animals of Kumamoto University and the Japan Neuroscience Society.

### Staining

Immunohistochemistry was conducted according to a standard protocol. Briefly, postnatal mouse brains were dissected and fixed with 4% paraformaldehyde (PFA) in PBS at room temperature (RT) for 3 h and then incubated sequentially with 12.5% and 25% (W/V) sucrose-containing PBS (pH 7.4), sequentially. After the brains were frozen at −80°C, coronal sections of 20 or 40 μm were cut with a cryostat and collected in PBS containing 0.1% sodium azide. For staining with anti-RORC and -NRN1 antibodies, sections were mounted on slide glasses and treated with 10 mM sodium citrate buffer (pH 6.0) at 105°C for 10 min or antigen retrieval solution (HistoVT One, Nacalai Tesque) at 70°C for 20 min, respectively. After blocking with PBS containing 5% normal goat/donkey serum and 0.1% Triton X-100 (blocking buffer) at RT for 1 h, the sections were incubated at 4°C overnight with the following antibodies in the blocking buffer: mouse anti-RORC (1:800; catalog no. PP-H3925-00, Perseus Proteomics; although this antibody recognizes RORα, β and γ, RORγ is not expressed in the postnatal brain, and RORα expression is weaker than and generally overlapping with RORβ expression in the cortex and thalamus (in Allen Brain Atlas)), rabbit anti-GFP (1:800; #A6455, Invitrogen), rabbit anti-Iba1 (1:500; catalog no. 019-19741, Wako), rabbit anti-5HTT (1:10000; catalog no. 24330, ImmunoStar), goat anti-Brn2 (1:50; catalog no. sc-6029, Santa Cruz), rat anti-Ctip2 (1:200; catalog no. ab18465, Abcam), rabbit anti-Tbr1 (1:500; catalog no. ab31940, Abcam), rabbit anti-RFP (1:1000; catalog no. PM005, MBL), rabbit anti-ssDNA (1:300; catalog no. 18731, MBL), rabbit anti-RORβ (1:5000; catalog no. pAb-RORβHS-100, Diagenode), mouse anti-NeuN (1:400; catalog no. MAB377, Chemicon), rabbit anti-NRN1 (1:50; catalog no. sc-25261, Santa Cruz), goat anti-VGF (1:50; catalog no. sc-10381, Santa Cruz). After three washes with PBS containing 0.1% Triton X-100 (PBST), the sections were incubated with Alexa 488- or Alexa 594-conjugated secondary antibody (1:500; Thermo Fisher Scientific) at RT for 2 h. For nonfluorescent detection, sections were incubated with biotinylated secondary antibodies and processed using the ABC histochemical method (VECTASTAIN ABC Kit, Vector). After three washes with PBST, the sections were counterstained with a DAPI solution (1:1000, Wako, Japan) and embedded with a mounting agent (SlowFade Gold antifade reagent, Thermo Fisher Scientific).

For EdU labeling of layer 4 neurons and postnatally generated cells, 5 mg/mL EdU in PBS was intraperitoneally injected into pregnant mutant mice (30 μg/g) at E14.7 (17:00) , and into perinatal ICR mice once a day for 4days (50 μg/g). Frozen sections prepared as described above were stained for EdU using a detection kit (Click-iT EdU Imaging Kit, Thermo Fisher Scientific). For Nissl staining, sections were immersed in 0.5% cresyl violet for several minutes and then subjected to an ethanol series (50%-75%-90%-100%) and embedded with a resin.

### Plasmids

For forced expression of NRN1 and VGF, pCAG-*Nrn1*-Flag and pCAG-*Vgf*-Fc, respectively, were used (Sato et al., 2012).

For electroporation-based cell ablation, pCAG-DTR was constructed as follows: a cDNA fragment containing the coding region of human HBEGF (GenBank accession number: BC033097) was obtained from a commercial supplier (clone ID 100067676, DNAFORM). The coding region was first cloned into a pGEM-T Eeasy vector cut with HincII and EcoRV followed by adenine addition and ligation. The coding region was subcloned into the pCAG vector by digesting pGEM-T Easy-DTR and pCAG with EcoRI and ligating them.

### In utero *electroporation*

*In utero* electroporation was carried out as described previously (Matsui et al., 2011; Saito and Nakatsuji, 2001; Tabata and Nakajima, 2001) with slight modifications. To exogenously express *Nrn1* and *Vgf* in cortical layer4, time-pregnant mutant mice of E14.3 were deeply anesthetized with pentobarbital (50 mg/kg). After the abdomen had been cleaned with 70% ethanol, a 3 cm midline laparotomy was performed, and the uterus was exposed. For DNA microinjection, plasmid DNA purified with the Plasmid Maxi-prep Kit (Genopure Plasmid Maxi Kit, Roche) was dissolved in Tris-EDTA buffer. Fast Green solution (0.1%) was added to the plasmid solution at 1:20 (v/v) ratio to monitor the injection. Approximately 1-2 μl of a mixture of 3 mg/ml pCAG-*Nrn1*-Flag and pCAG-*Vgf*-Fc and 1 mg/ml pCAG-DsRed was injected into the lateral ventricle with a glass micropipette. The embryos in the uterus were placed in a tweezers-type electrode equipped with two platinum discs of 5 mm in diameter at the tip (CUY650-5, Nepa Gene, Japan). Electronic pulses (30 V, 50 ms) were delivered four times at intervals of 950 ms with an electroporator (NEPA21, Nepa Gene, Japan), and then, the uterine horns were placed back into the abdominal cavity. The abdominal wall and skin were sewed up with surgical sutures, and the embryos were allowed to develop until P7. For electroporation-based cell ablation of thalamic neurons, time-pregnant ICR mice of E11.5 were used and a mixture of 3 mg/ml pCAG-DTR and 1 mg/ml pCAG-DsRed was injected into the third ventricle, followed by application of electric pulses (25 V, 50 ms).

### *Generation of* Nrn1 *and* Vgf-*deficient mice by the CRISPR/Cas9 method*

To delete *Nrn1* and *Vgf* loci, one and two sgRNAs targeting the *Nrn1* and *Vgf* exons, respectively, were designed (*Nrn1* sgRNA: AGCATGGCCAACTACCCGCA; *Vgf* sgRNA: TCACGTTGCCGGCATCCGTC, CGGTACTGTTGCAGGCACTGGACCGT). Fertilized eggs derived from C57BL/6J mice were electroporated with a mixture of sgRNAs and Cas9 protein and transplanted into foster mother mice. Brains were collected from pups at P8 and fixed with 4% PFA in PBS for at RT 3 h. Before fixation, a piece of cerebellum was dissected. Genomic DNA was extracted and *Nrn1* and *Vgf* loci were analyzed by sequencing of. For genomic sequencing, DNA fragments including the sgRNA sequences were amplified by PCR using the following primers: *Nrn1*: 5’-ACCAGGGAACTGAGCCTGAG-3’ and 5’-GGACTCACCTCCCTGCTATC-3’; *Vgf*: 5’-GGTACCCAGAAGGAGGATTG-3’ and 5’-TTGCTCGGACTGAAATCTCG-3’. Sequencing PCR was performed using the amplified DNA fragments as a template and the primers: *Nrn1*: 5’-ACCAGGGAACTGAGCCTGAG-3’; *Vgf*: 5’-GGTACCCAGAAGGAGGATTG-3’ (near sgRNA#1) or 5’-CTCAGCTCTGAGCATAATGG-3’ (near sgRNA#2). Mice harboring a deletion that causes frame shift resulting in translation failure without wild-type sequences nor deletion in multiples of three bases resulting in truncated protein product were designated null mutants.

### Lipophilic dye labeling

For labeling neural connections, P7 mutant mouse brains were fixed with 4% PFA in PBS at RT for 3 h. A small crystal of DiA and DiI (catalog no. D3883 for DiA and D3911 for DiI, Thermo Fisher Scientific) was inserted into S1 and V1, respectively. Incubated in 4% PFA in PBS at 37°C for two weeks after implantation, the brains were cut into 100 μm slices with a vibratome (Leica) and observed by fluorescence microscopy (BX52, Olympus).

### Data quantification and statistical analysis

Marker-positive or EdU-labeled cells were counted in photomicrographs of single focal plane (850 × 850 μm) acquired with a laser scanning confocal microscope (LSM780, Zeiss) with a 10x objective lens. Particles of >13.37 μm^2^ with signal intensities higher than a given threshold, were quantified using Metamorph software (Molecular Devices). The relative cell number and SEM were calculated for each experimental condition. To measure the thickness of the marker-positive layer, the maximum radial distance of the tangential lines, along which more than two positive particles defined as above were distributed, was measured automatically by Metamorph. The relative thickness and SEM were calculated for each experimental condition. To measure the distribution variation of layer 4 cells in the barrel field, the region of 25 μm in height, and 583 μm in width, was extracted from confocal images of the barrel field stained with DAPI. The region was subdivided into 35 (25 × 16.7 μm each) and total intensity was measured for each subdivision. The variation of intensities among 35 subdivisions was calculated for each region. Group means were compared using Student’s *t*-test, and \**P*<0.05 was regarded statistically significant.

