## Supplementary material for "Thalamocortical axons control the cytoarchitecture of neocortical layers by area-specific supply of secretory proteins": Figure-figure supplement

**Figure Supplements with Legends**

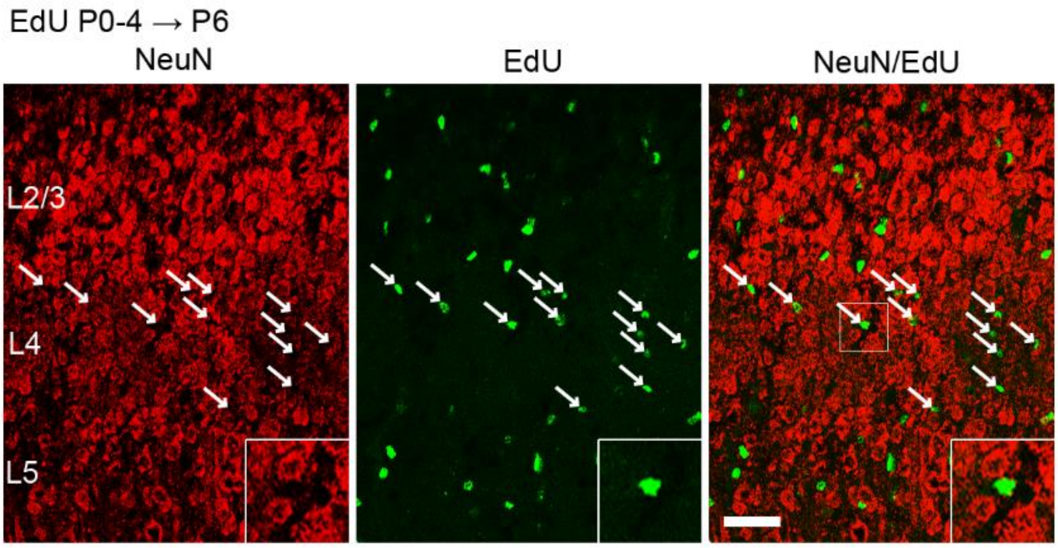


**Figure 1-Figure supplement 1.** Layer 4 neurons are not additively generated postnatally**.**

Coronal sections of the S1 cortex of a P6 mouse administrated EdU every day from P0 to P4 to label newly generated postmitotic cells, double stained for NeuN and EdU. Note that all the EdU-labeled cells in layer 4 (L4) were not co-stained for NeuN (arrows), indicating that they are not neurons. Bottom-right insertions are magnified view of square region in the merged image. L2/3, layer 2 and 3; L5, layer 5. Scale bar, 50 μm.

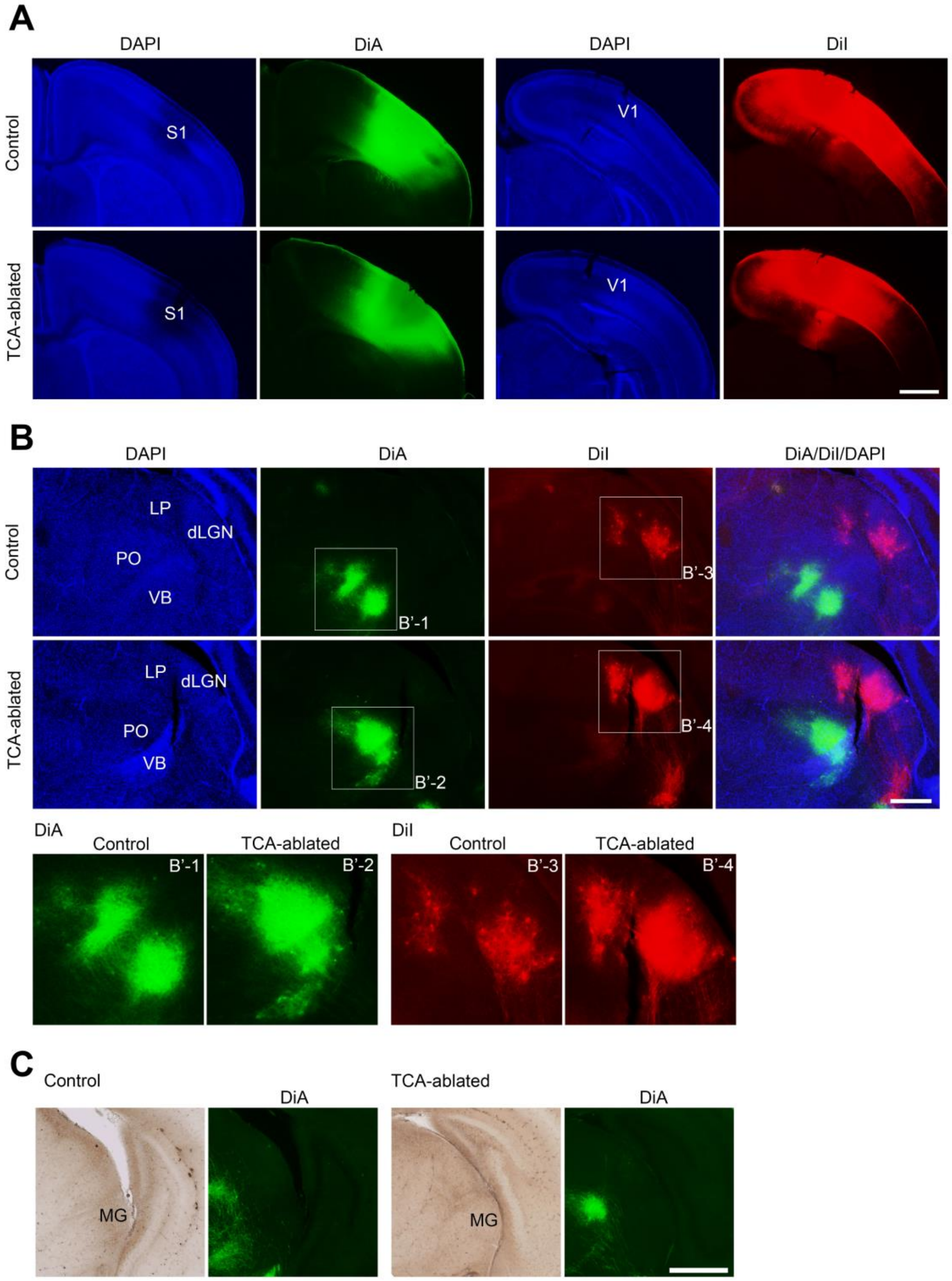


**Figure 2-Figure supplement 1.** TCA connections in S1 of TCA-ablated mouse.

(**A**) Coronal sections of control and TCA-ablated cortices showing DiA (green) and DiI (red) embedded in S1 and V1, respectively. (**B, C**) Coronal sections of thalami of control and TCA-ablated mice showing retrogradely labeled cell bodies and anterogradely labeled axons from the cortex in the thalamus. Note that DiA signal was not observed in dLGN nor MG in the TCA-ablated mouse. Panels B'-1-4 are magnified view of squares in b. dLGN, dorsal lateral geniculate nucleus; LP, lateral posterior nucleus; MG, medial geniculate nucleus; PO, posterior nucleus; S1, primary somatosensory area; VB, ventrobasal nucleus; V1, primary visual area. Scale bars, (**A**) 1 mm, (**B**) (**C**) 500 μm.

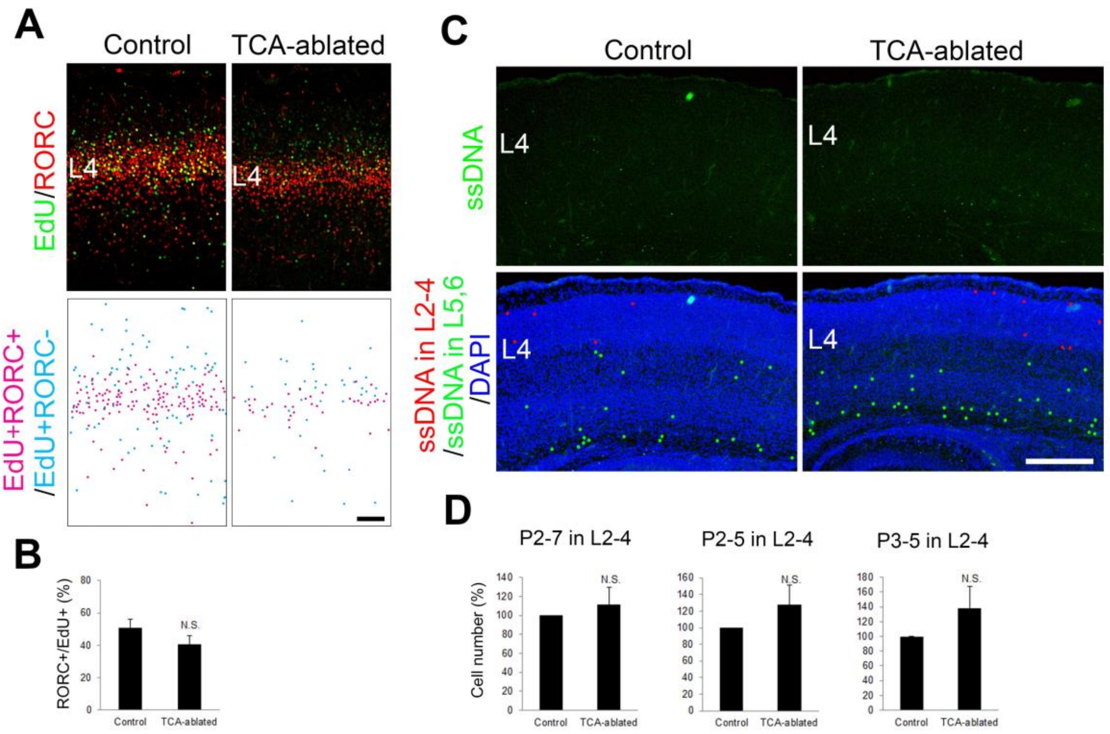


**Figure 3-Figure supplement 1.** Effects of TCA ablation on cell fate and cell death of layer 4 neurons in S1.

(**A**) Cross sections of S1 cortex of control and TCA-ablated mice to which EdU had been administrated at E14.7, stained for EdU and RORβ. Cells double positive for EdU and RORβ and cells single positive for EdU, but negative for RORβ, are represented as magenta dots and cyan dots (labeled manually), respectively, in the lower panels. (**B**) Quantification of the results. The percentage of RORβ-expressing cells among EdU-labeled cells is presented as the mean ± SEM: control, 50.94 ± 5.05%; TCA-ablated, 40.82 ± 5.07%; *N* = 6 sections from 3 mice for each, *P*=0.115; *t*-test. Note that the proportion of RORβ-positive cells within EdU-labeled cells was not significantly affected in TCA-ablated mice. (**C**) Coronal sections of S1 cortex of control and TCA-ablated mice at P4, stained for ssDNA. Stained cells in layers 2-4 (L2-4) and layers 5-6 are represented as red and green dots, respectively, in the DAPI-stained image. (**D**) Quantification of the results. The number of ssDNA-positive cells in layers 2-4 was not significantly increased in TCA-ablated mice. Summation data of P2-7, P2-5, and P3-5 mice are shown as a percentage of the control (mean ± SEM): P2-7, 111.69 ± 18.18%, $N$ = 8 sections from 7 mice, $P$=0.274; P2-5, 127.59 ± 23.65%, $N$ = 6 sections from 4 mice, $P$=0.163; P3-5, 138.41 ± 29.74%, $N$ = 4 sections from 3 mice, $P$=0.162; *t*-test. The same number of control animals was used. L4, layer 4. Scale bars, (**A**) 100 µm, (**C**) 500 µm.

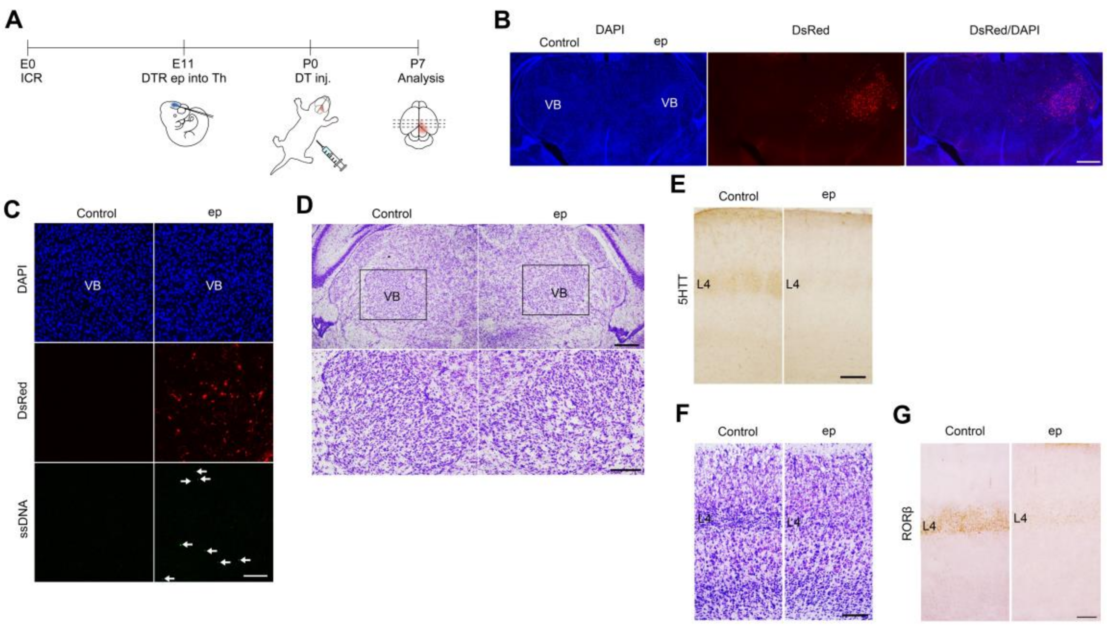


**Figure 3-Figure supplement 2.** Electroporation-based TCA ablation causes reduction in the number of layer 4 neurons.

(**A**) Schematic representation of the experimental procedure used for electroporation-mediated thalamic ablation. DTR-encoding plasmid together with DsRed plasmid was electroporated into the thalamus of E11.5 ICR mice. DT was administrated at P0 and brains were collected at P7. (**B-D**) Coronal sections of thalamus specimens of electroporated mice at P7 (**B**, **D**) and P2 (**C**). VB neurons were preferentially electroporated, as shown by DsRed fluorescence on the electroporated (ep) side of the thalamus at P7. Upon administration of DT at P0, a number of ssDNA-positive dying cells (arrows in **C**) were detected on the ep side, where residual DsRed-expressing cells are visible at P2. Nissl staining revealed a reduction in neuron in the VB on the ep side at P7. (**E-G**) Coronal sections of S1 cortex of a DTR-electroporated specimen to which DT was administrated at P0. 5-HTT immunoreactivity was markedly decreased in layer 4 on the ep side (**E**). Nissl staining and RORC immunohistochemistry revealed reduction in cell density (**F**) and RORβ-expressing cells (**G**) in layer 4 on the ep side. VB, ventrobasal nucleus. Scale bars, (**B**) 2 mm, (**C**) 500 µm, (**D**) (**E**) (**F**) (**G**) 200 µm.

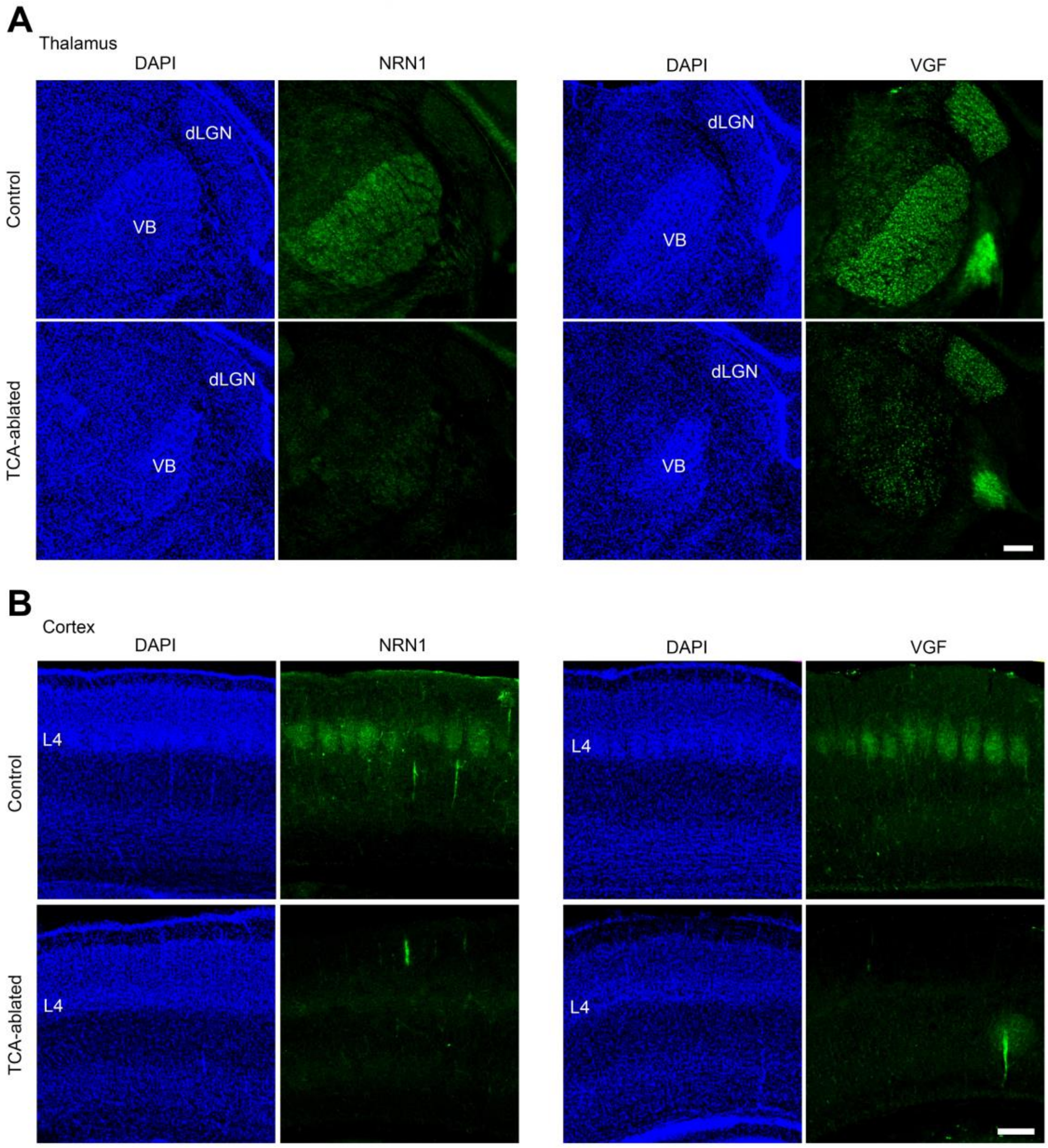


**Figure 5-Figure supplement 1.** Expression of NRN1 and VGF are lost in the cortex of TCA-ablated mice.

Coronal sections of (**A**) thalamus and (**B**) S1 cortex of control and TCA-ablated mice at P7 immunostained for NRN1 and VGF. Note that NRN1 and VGF are expressed in the VB of control, but not TCA-ablated mice, and that their signals are absent in layer 4 of S1 cortex in TCA-ablated mice. dLGN, dorsal lateral geniculate nucleus; L4, layer 4; VB, ventrobasal nucleus. Scale bars, 200μm.

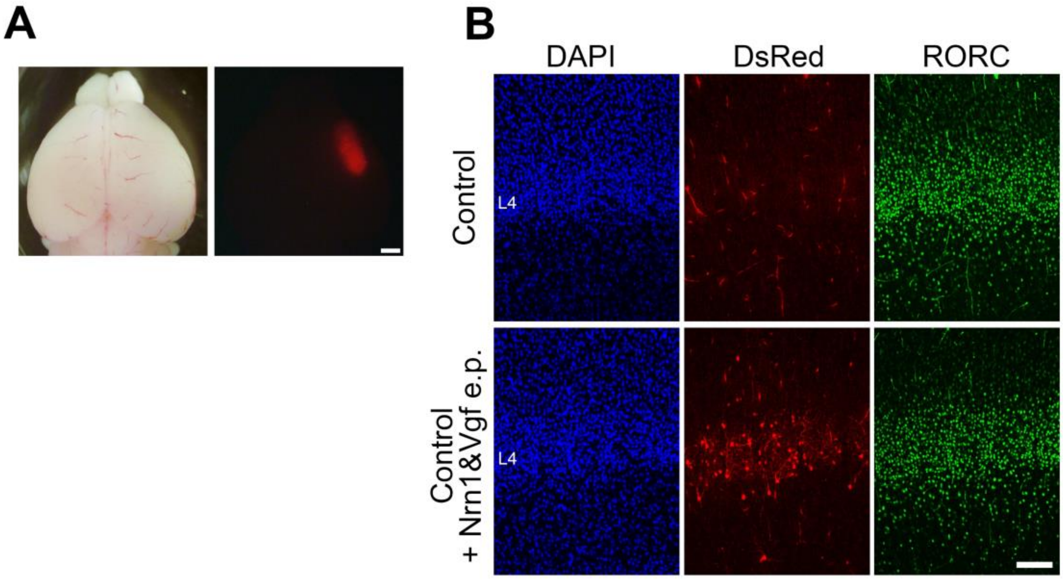


**Figure 5-Figure supplement 2.** Overexpression of NRN1 and VGF did not affect layer 4 formation in control mice.

(**A**) P7 electroporated brain showing DsRed signal in the right hemisphere. (**B**) Cross sections of S1 cortices of control and control + electroporated (e.p.) mice stained for DsRed and RORβ. Scale bars, (**A**) 1 mm, (**B**) 100 µm.

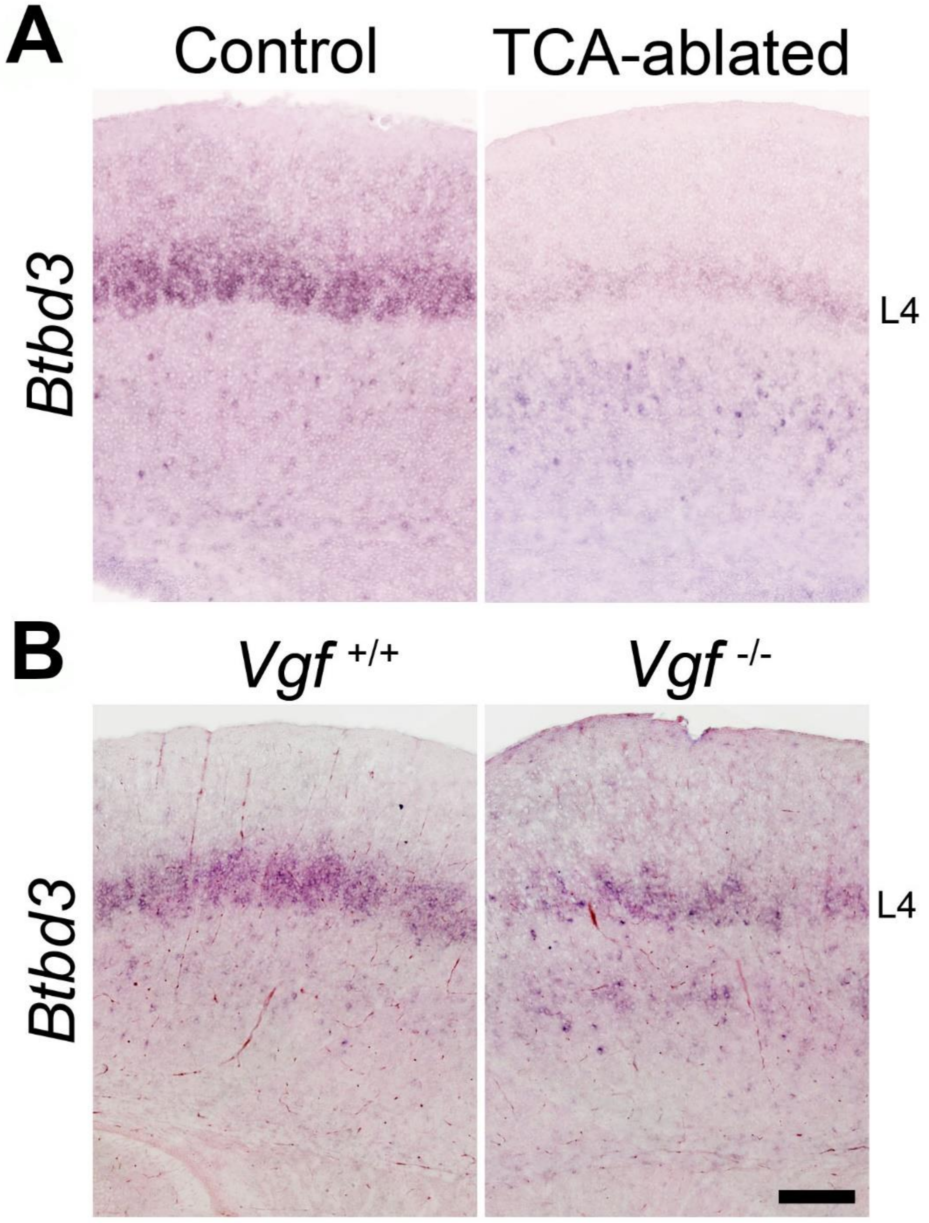


**Figure 7-Figure supplement 1.** Expression of an activity-dependent gene *Btbd3* in layer 4 neurons.

In situ hybridization for *Btbd3* in S1 layer 4 of P7 mice. (**A**) TCA-ablated and (**B**) *Vgf*-KO mice with controls. L4, layer 4. Scale bar, 100 µm.

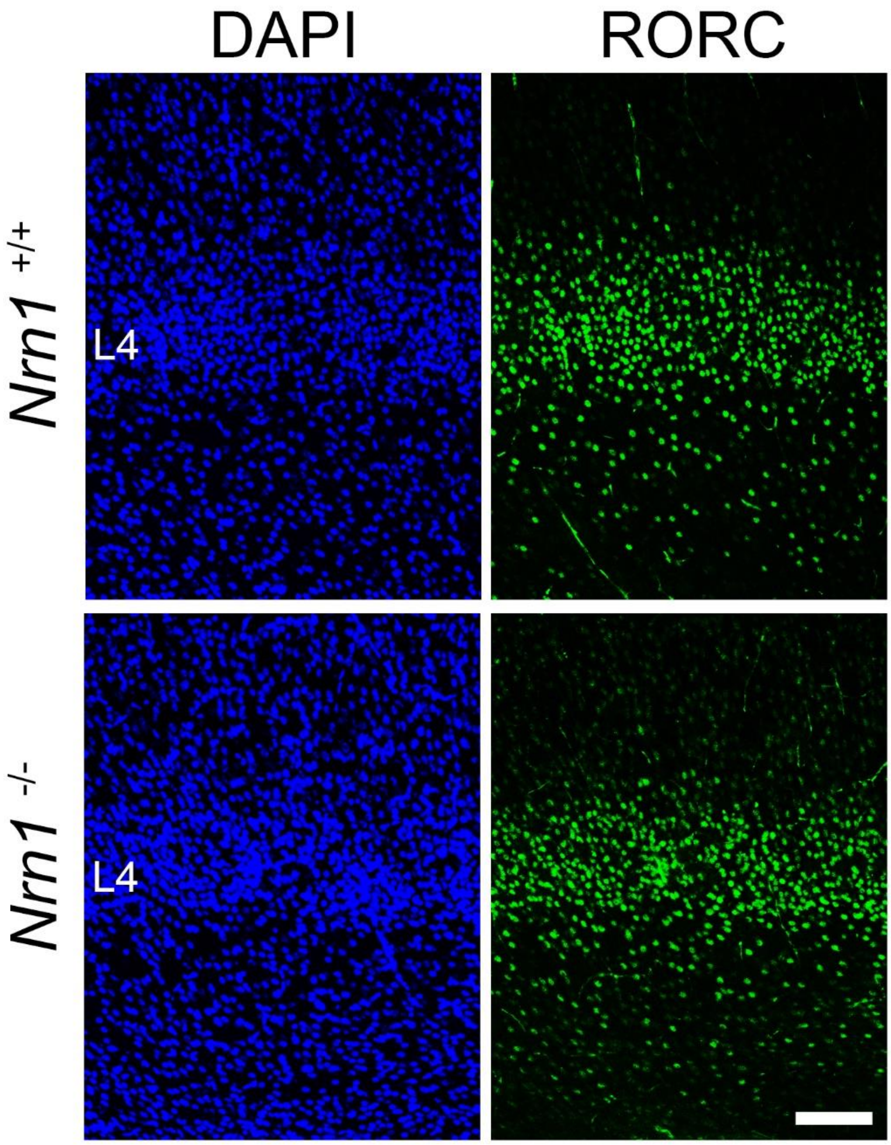


**Figure 7-Figure supplement 2.** *Nrn1*-KO mice show normal layer 4.

Coronal sections of S1 cortex of wild-type and *Nrn1*-deficient mice at P8 stained with anti-RORC antibody. Scale bar, 100 μm.
